# ScHiCAtt: Enhancing Single-Cell Hi-C Resolution Using Attention-Based Models

**DOI:** 10.1101/2024.12.16.628505

**Authors:** Rohit Menon, H. M. A. Mohit Chowdhury, Oluwatosin Oluwadare

**Affiliations:** Department of Computer Science, University of Colorado at Colorado Springs, Colorado Springs, 80918, USA; Department of Biomedical Informatics, University of Colorado Anschutz Medical Campus, Aurora, 80045, USA

**Keywords:** Hi-C data, Self-Attention, Resolution Enhancement, Single-cell Hi-C, Data Sparsity

## Abstract

The spatial organization of chromatin is fundamental to gene regulation and essential for proper cellular function. The Hi-C technique remains the leading method for unraveling 3D genome structures; however, limited resolution, data sparsity, and incomplete coverage in single-cell Hi-C data pose significant challenges for comprehensive analysis. Traditional CNN-based models often suffer from blurring and loss of fine details, while GAN-based methods encounter difficulties in maintaining diversity and generalization. Moreover, existing algorithms perform poorly in cross-cell line generalization, where a model trained on one cell type is used to enhance high-resolution data in another cell type. To address these limitations, we propose ScHiCAtt (Single-cell Hi-C Attention-Based Model), which leverages attention mechanisms to capture both long-range and local dependencies in Hi-C data, significantly enhancing resolution while preserving biologically meaningful interactions. We implement this mechanism and check its validity on data from different cells of the same organisms and data of different organisms. By dynamically focusing on regions of interest, attention mechanisms effectively mitigate data sparsity and enhance model performance in low-resolution contexts. Extensive experiments on Human and Drosophila single-cell Hi-C data demonstrate that ScHiCAtt consistently outperforms existing methods in terms of computational and biological reproducibility metrics across different downsampling ratios, especially under extreme downsampling conditions. The model is publicly available at https://github.com/OluwadareLab/ScHiCAtt.

## 1 Introduction

Three-dimensional (3D) conformation of chromosomes is crucial for elucidating genomic processes within the nuclei of eukaryotic cells. The Hi-C technique facilitates an all-versus-all mapping of chromosomal fragment interactions, resulting in an interaction frequency contact matrix, where *n×n* represents the number of fragments in a chromosome or genome at a specific resolution, Lieberman-Aiden et al., 2009. These Hi-C data are critical for numerous algorithms designed to improve the understanding of genome organization, Oluwadare et al., 2019. A major challenge in this field is the scarcity of high-resolution Hi-C data, which are indispensable for identifying intricate genomic topologies such as enhancer-promoter interactions and subdomains.

To address this need, deep learning models have been employed to predict high-resolution data from low-resolution data with remarkable accuracy. Notable models in this area include HiCPlus Y. Zhang et al., 2018, HiCNN T. Liu and Z. Wang, 2019a, hicGAN Q. Liu et al., 2019, Boost-HiC Carron et al., 2019, HiCSR Dimmick, 2020, SRHiC Z. Li and Dai, 2020, HiCNN2 T. Liu and Z. Wang, 2019b, HiCARN Hicks and Oluwadare, 2022, and DeepHiC Hong et al., 2020. These models leverage various network architectures such as Convolutional Neural Networks (CNNs), Autoencoders, and Generative Adversarial Networks (GANs). Despite the advancements made by these models, there remains considerable room for improvement, especially when it comes to single-cell Hi-C data enhancement, Y. Wang et al., 2023, as all of the aforementioned methods are designed for bulk Hi-C data enhancement.

Single-cell Hi-C (scHi-C) is a groundbreaking technology that offers a unique opportunity to investigate 3D genome structures at the single-cell level with high resolution, Galitsyna and Gelfand, 2021. By capturing chromatin interactions at the individual cell level, scHi-C enables the exploration of cellular heterogeneity in chromatin conformation, Arrastia et al., 2020; Collombet et al., 2020; Payne et al., 2021. However, scHi-C data are characterized by high dimensionality, noise, and sparsity, presenting computational challenges that demand innovative solutions for the accurate reconstruction of 3D genome structures, Paulsen et al., 2015; Galitsyna and Gelfand, 2021. Therefore, scHi-C data imputation is crucial, as it enables the reconstruction of enhanced contact maps from raw and sparse scHi-C data, thereby improving the quality for downstream analyses, including the reconstruction of chromatin organization at the single-cell level. This enhancement aids in uncovering cell-to-cell variability and heterogeneity, ultimately providing deeper insights into cellular functions and disease mechanisms Y. Wang et al., 2023.

Recently, algorithms like ScHiCEDRN, Y. Wang et al., 2023 and Loopenhance, S. Zhang et al., 2022 have been developed to address the challenges of scHi-C data enhancement. While these methods aim to improve the resolution of single-cell Hi-C data, they often fall short in capturing the complex spatial relationships within chromatin structures, especially long-range dependencies. This limitation leads to the loss of critical interactions, which are essential for accurately reconstructing chromatin topology.

On the other hand, Attention mechanisms have proven effective in capturing both short-range and long-range dependencies in various domains, such as natural language processing and computer vision, Vaswani, 2017. These mechanisms enable models to focus on different regions of the input data dynamically; hence, they have the potential to be used to enhance the resolution of sparse datasets like scHi-C by capturing context at multiple scales. The motivation behind our work is to leverage Attention mechanisms to address challenges unique to scHi-C data, such as sparsity, noise, and limited coverage. By selectively focusing on relevant chromatin interactions, our approach aims to provide a more biologically meaningful reconstruction of 3D genome structures.

In this work, we propose ScHiCAtt, which employs a cascading residual network integrated with an optimal attention mechanism identified through validation across multiple candidates. ScHiCAtt explores different attention mechanisms, such as self-attention, local attention, global attention, and dynamic attention (Attention-in-Attention), selecting the optimal mechanism for each layer during training to determine the best attention mechanism to incorporate for scHi-C data enhancement. The goal of this experimentation is to allow ScHiCAtt to capture both short-range and long-range dependencies adaptively, thus enhancing the quality of scHi-C data reconstruction.

Through comprehensive experiments on human and Drosophila data across various downsampling rates, we demonstrate that ScHiCAtt significantly improves the resolution of scHi-C data. Our results show superior performance in terms of computational metrics and biological reproducibility metrics, such as GenomeDISCO, Ursu et al., 2018, compared to existing methods, particularly under extreme downsampling conditions. Moreover, ScHiCAtt maintains efficient training times, making it a robust solution for high-resolution single-cell Hi-C data enhancement.

## 2 Materials and Methods

### 2.1 Model Architecture

Our model architecture starts with an entry convolution layer (Figure 1A) that processes the input raw scHi-C contact map. This is followed by a series of cascading blocks interleaved with attention layers, designed to progressively upscale the resolution of the Hi-C maps. The final high-resolution Hi-C maps are produced through an exit convolution layer. The architecture also includes tunable hyperparameters such as the number of cascading blocks and attention layers, allowing for flexibility in optimizing the model’s performance.

**Figure 1.**
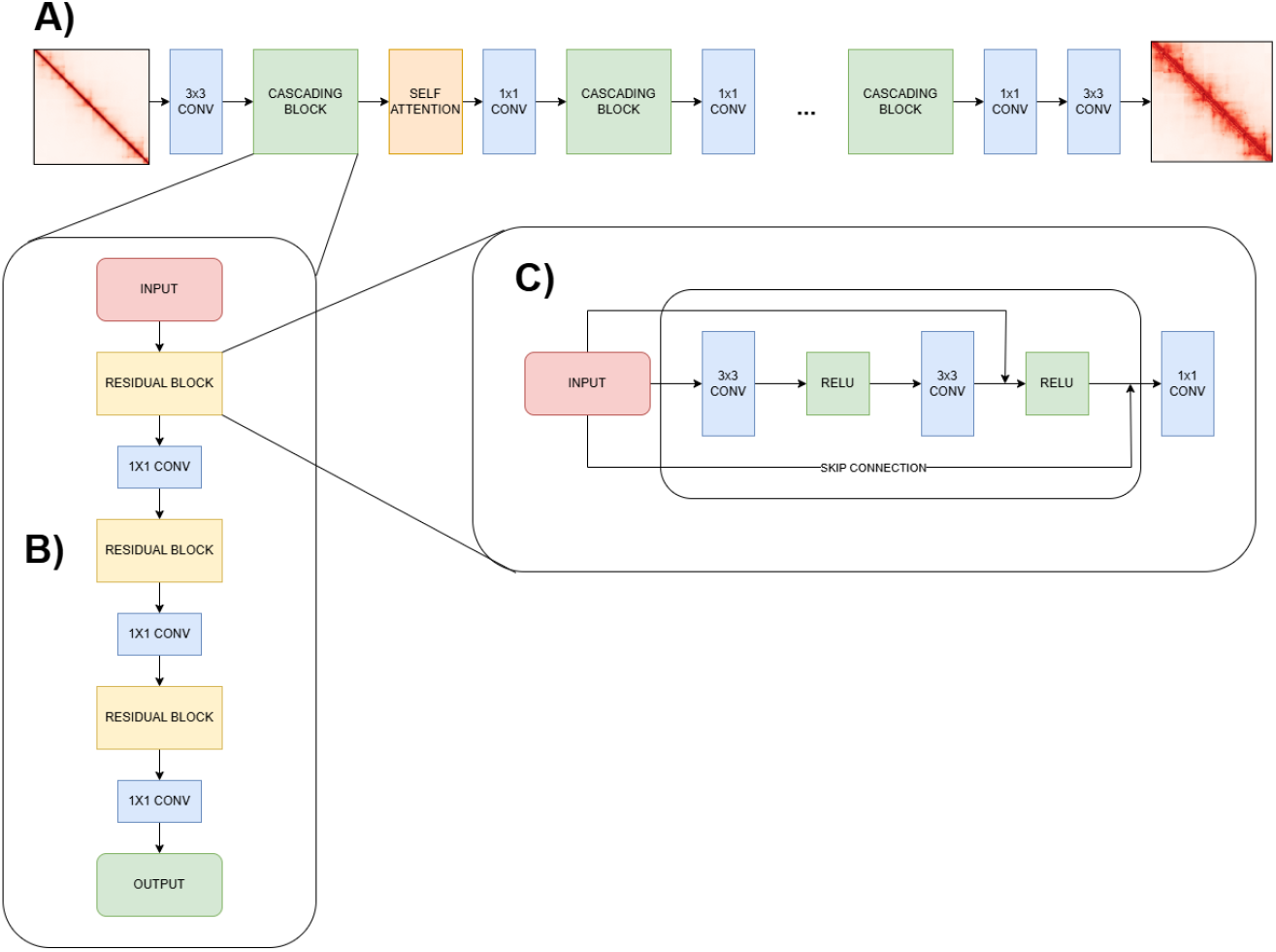
Architecture of the Cascading Residual Network with Attention for Hi-C Super Resolution. **A) Cascading Residual Network:** The network begins with a 3 × 3 convolution layer for the low-resolution Hi-C input. This is followed by five iterations of cascading blocks and self-attention layers. Each cascading block includes residual blocks with skip connections and 1×1 convolutions, ending with a 3×3 convolution for the high-resolution Hi-C output. **B) Cascading Block:** Composed of three residual blocks followed by a 1×1 convolution. Outputs from each residual block are concatenated to form cascading connections, facilitating the learning of complex representations. **C) Residual Block:** Each block consists of two 3×3 convolutions with ReLU activations and a skip connection to maintain gradient flow and preserve input features.

In the following subsections, we explore various attention mechanisms that have been considered in our study. We describe each mechanism in detail, highlighting its unique features and the rationale behind its selection for our research. Furthermore, we elucidate how these mechanisms were implemented within our architecture for evaluation.

#### 2.1.1 Self-Attention Mechanism

The self-attention mechanism in our architecture (Figure 1A) facilitates efficient learning of both local and global chromatin interactions by allowing the model to dynamically assign weights to relationships between chromatin loci, regardless of their spatial distance on the Hi-C contact maps. This capability is crucial for capturing both short-range and long-range dependencies within chromatin structures.

To achieve this, the attention scores are computed by taking the scaled dot-product between the input projection matrices: queries (**H**) and keys (**J**), divided by the square root of the keys dimension. The resulting attention scores are passed through a softmax function to compute the attention weights, which are then applied to the values (**Z**). This enables the model to prioritize important interactions, enhancing the quality of the predicted high-resolution contact maps.

The process is defined as:

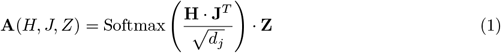

Here, **H** ∈ ℝ^*n*×*d*^, **J** ∈ ℝ^*n*×*d*^, and **Z** ∈ ℝ^*n*×*d*^ represent the query, key, and value matrices respectively, where *n* is the sequence length (number of loci in the Hi-C contact map), and *d* is the feature dimension. The term *d*_*j*_ is the dimension of the keys (i.e., *d*_*j*_ = *d*) used to scale the dot product and stabilize the training process. This mechanism enables the model to focus on critical chromatin interactions, significantly improving prediction accuracy.

#### 2.1.2 Cascading Residual Blocks

The backbone of our architecture is the cascading residual blocks, illustrated in Figure 1B, Ahn et al., 2018. Each block comprises residual units with skip connections that progressively refine the Hi-C contact maps. These cascading blocks are interconnected, allowing for the aggregation of features across different layers.

#### 2.1.3 Local Attention Mechanism

Local attention is applied within the cascading residual blocks (Figure 1B). It focuses on capturing fine-grained chromatin interactions within localized regions of the Hi-C contact maps. The use of depthwise and pointwise convolutions in the local attention mechanism allows the model to enhance the spatial resolution of the Hi-C maps by emphasizing intricate local details.

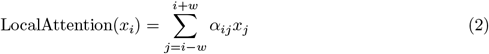

where 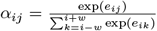, and *e*_*ij*_ = (*x*_*i*_*W*_*Q*_)(*x*_*j*_*W*_*K*_)^*T*^. Here, *x*_*i*_ is the input at position *i, w* is the window size defining the local neighborhood, *W*_*Q*_ and *W*_*K*_ are learnable weight matrices for queries and keys respectively.

#### 2.1.4 Global Attention Mechanism

The global attention mechanism is applied after several cascading residual blocks (Figure 1B) to ensure that global chromatin structures are preserved. This module aggregates context across the entire Hi-C map and allows the model to capture large-scale genomic interactions, which are critical for accurate super-resolution Zhu et al., 2021.

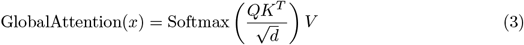

where *Q* = *xW*_*Q*_, *K* = *xW*_*K*_, *V* = *xW*_*V*_, and *W*_*Q*_, *W*_*K*_, *W*_*V*_ are learnable weight matrices for queries, keys, and values, respectively.

#### 2.1.5 Multi-Head Attention Mechanism

The multi-head attention mechanism is designed to enhance the model’s ability to capture complex relationships in the input data by dividing the input into multiple attention heads. Each head performs attention operations independently, focusing on different aspects of the input, which allows the model to extract diverse contextual information.

The mechanism takes three primary inputs: the Query (*Q*), Key (*K*), and Value (*V*) matrices. These inputs are derived from the original data through linear transformations. The attention operation for each head calculates a weighted representation of the Value matrix, where the weights are determined by the similarity between the Query and Key matrices. This is expressed mathematically as:

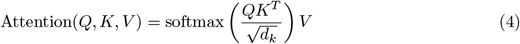

Here, *d*_*k*_ is the dimensionality of the Key matrix, and the softmax function ensures that the weights sum to 1, highlighting the most relevant features for each Query.

For multi-head attention, the inputs are split into *h* separate heads, each with its own *Q, K*, and *V*. The outputs from all heads are concatenated and passed through a final linear transformation, as shown in the equation below:

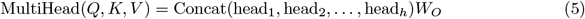

In this equation, head_*i*_ represents the output of the *i*-th attention head, and *W*_*O*_ is the learned weight matrix for the final linear transformation. This design allows the model to integrate information from multiple perspectives, improving its ability to capture chromatin interactions and other complex patterns.

#### 2.1.6 Dynamic Attention Mechanism

Dynamic Attention, also referred to as the Attention-in-Attention (A2A) mechanism, combines static and dynamic attention features to weigh their contributions adaptively. The dynamic attention module applies global pooling, followed by fully connected layers, to dynamically adjust the contribution of features based on and without attention Huang et al., 2019.

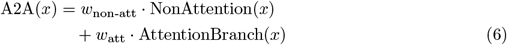

### 2.2 Loss Function

To optimize the quality of the enhanced scHi-C contact matrices, we leverage several key loss functions that address distinct aspects of the reconstruction process. These loss functions ensure that the generated matrices not only minimize pixel-wise error with respect to the target but also maintain structural integrity and visual consistency.

#### 2.2.1 Mean Squared Error (MSE)

The goal is to minimize the pixel-wise difference between the true and enhanced scHi-C matrices, ensuring that the generated maps closely approximate the true scHi-C data.

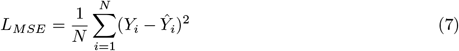

In this equation:

- *N* : The total number of data points or pixels in the scHi-C matrices.
- *Y*_*i*_: The true value of the *i*-th pixel in the scHi-C matrix.
- *Ŷ*_*i*_: The predicted value of the *i*-th pixel in the enhanced scHi-C matrix.
- *L*_*MSE*_: The computed Mean Squared Error, representing the average of the squared differences between the true and predicted values.

This loss function penalizes larger deviations more heavily due to the squaring operation, encouraging the model to generate outputs that closely match the true data.

#### 2.2.2 Perceptual Loss

Perceptual loss, based on feature representations from a pre-trained VGG network Wu et al., 2020, ensures that the generated Hi-C maps are not only pixel-accurate but also visually consistent with the real Hi-C data.

In the perceptual loss *L*_*V GG*_, we utilize the feature maps from specific layers of the pre-trained VGG network:

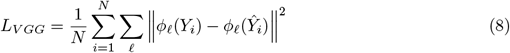

where *ϕ*_*𝓁*_(*·*) denotes the feature map extracted from the *𝓁*-th layer of the VGG network.

#### 2.2.3 Total Variation (TV) Loss

TV loss reduces noise and enforces smoothness in the generated Hi-C maps, improving the overall visual quality.

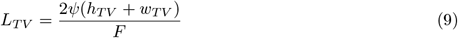

#### 2.2.4 Adversarial Loss (AD)

Adversarial loss improves the realism of the generated high-resolution Hi-C maps by ensuring that the discriminator cannot easily distinguish between real and generated matrices.

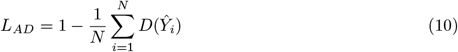

### 2.3 Evaluation Metrics

To evaluate the effectiveness of our models in enhancing the resolution of scHi-C data, we used a few standard metrics that give us different ways to look at the quality of the reconstructed contact maps. Each of these metrics helps us understand how good the reconstruction is from different perspectives. They can broadly be categorized as computational metrics, such as Structural Similarity Index Measure, Peak Signal-to-Noise Ratio, and Signal-to-Noise Ratio and biological reproducibility metrics, such as GenomeDISCO, Ursu et al., 2018.

#### 2.3.1 Structural Similarity Index

Structural Similarity Index Measure(SSIM) quantifies the structural similarities between the true and enhanced scHi-C matrices.

SSIM is defined as,

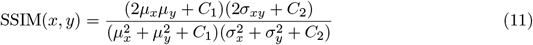

Here, *µ*_*x*_ and *µ*_*y*_ are the means of *x* and *y*, 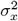 and 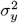 are the variances, *σ*_*xy*_ is the covariance between *x* and *y*, and *C*_1_ and *C*_2_ are constants to stabilize the division when the denominator is close to zero.

### 2.3.2 Peak Signal-to-Noise Ratio

As the name states, Peak Signal-to-Noise Ratio (PSNR) quantifies the ratio between the maximum achievable signal and the noise that distorts it.

PSNR is defined as

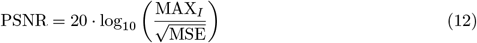

In this equation:

- PSNR: Peak Signal-to-Noise Ratio, a metric to measure the quality of the enhanced image.
- MAX_*I*_ : The maximum possible pixel value of the image (e.g., 255 for 8-bit images).
- MSE: Mean Squared Error between the original and enhanced images.
- log_10_: The base-10 logarithm.

#### 2.3.3 Mean Squared Error

Mean Squared Error (MSE) calculates the average squared difference between the predicted and true values.

Mean Squared Error is defined as,

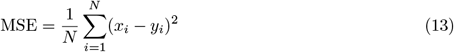

#### 2.3.4 Signal-to-Noise Ratio

Signal-to-Noise Ratio (SNR) measures the relationship of the signal power to noise power.

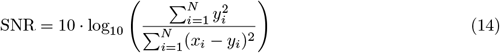

#### 2.3.5 GenomeDISCO

In this study, we utilize GenomeDISCO Ursu et al., 2018 as a measure of biological reproducibility. GenomeDISCO produces a concordance score ranging from −1 to 1, reflecting the biological similarity between two contact maps. A higher value indicates better concordance. The methodology entails applying a smoothing technique to the contact maps through their graph representations, followed by the calculation of the similarity score on the resulting smoothed matrices.

## 3 Results

### 3.1 Dataset Preparation

For this study, we utilized scHi-C datasets as prepared by the ScHiCEDRN framework, which includes data from both *Drosophila melanogaster* and *Homo sapiens* cell lines. The *Drosophila* dataset comprises seven chromosomes (chr2L, chr2R, chr3L, chr3R, chr4, chrX, and chrM) (GSE13181 Ulianov et al., 2021, while the human dataset includes chromosomes from the frontal cortex (GSE130711) Lee et al., 2019; Luo et al., 2022.

Following the preprocessing steps as described in the ScHiCEDRN framework Y. Wang et al., 2023, we utilized the low-resolution contact maps provided, which had been downsampled to varying degrees (75%, 45%, 10% and 2% of the original raw reads). Detailed preprocessing information can be found in ScHiCEDRN, Y. Wang et al., 2023, and the datasets are publicly available at https://github.com/BioinfoMachineLearning/ScHiCEDRN. No additional preprocessing was performed on the data. For the human cell line, chromosomes 1, 3, 5, 7, 8, 9, 11, 13, 15, 16, 17, 19, 21, and 22 from Human cell 1 were used as the training dataset, while chromosomes 4, 14, 18, and 20 were used for validation. For testing, we used chromosomes 2, 6, 10, and 12 from both Human cell 1 and a different human cell, referred to as Human cell 2, as done by ScHiCEDRN. For testing on Drosophila cells, we used chromosomes chr2L and chrX.

These datasets were used as inputs for our models, with the raw scHi-C contact maps serving as the ground truth for model training and evaluation.

### 3.2 Hyperparameter Search for Individual Attention Mechanisms

We have conducted an extensive hyperparameter search to determine the optimal configuration for our architecture. The two criteria to optimize are (i) determining the best-performing attention mechanism and (ii) its placement within the network layers. Our primary focus is on the implementation of various attention mechanisms, including Self-Attention, Local Attention, Global Attention, and Dynamic Attention. The goal is to ascertain which attention mechanism and its placement within the network layers yield the best performance metrics, specifically the PSNR, SSIM, and SNR metrics. The loss function applied in this search is the MSE loss.

We performed experiments on the Human cell 1 dataset by integrating each attention mechanism in different layers of the model (Layers 2, 3, and 5) and evaluated their impact on the model’s performance. The average results, which were obtained from the corresponding validation set chromosomes are in Figure 2, Supplementary Figure S1 and Table I, indicate that the choice of attention mechanism and its placement within the network significantly influences the model’s output quality. As illustrated in Table I, the model configuration that utilized Self-Attention at Layer 2 consistently outperformed the other configurations across all metrics. Implying that the Layer 2 effectively captures both local and global chromatin interactions, enhancing the model’s ability to preserve long-range dependencies while refining the details in the contact maps. Specifically, it achieved high values of PSNR, SSIM and SNR (Table I). The Dynamic Attention mechanism at the same layer closely followed these results. Conversely, the Local and Global Attention mechanisms, while still providing significant improvements over a baseline model, did not achieve the same level of performance.

**Table I:** Comparison of each Attention Mechanism Performance at Different Layers using evaluation metrics: PSNR, SSIM, SNR, and GenomeDISCO on the Human cell 1 dataset.

| Attention Mechanism | PSNR | SSIM | MSE | SNR | GenomeDISCO | Layer |
| --- | --- | --- | --- | --- | --- | --- |
| Self-Attention | **39.77** | **0.9823** | 0.0010 | **5533.887** | **0.9181** | 2 |
| Local Attention | 32.24 | 0.9124 | 0.0015 | 5072.432 | 0.8454 | 2 |
| Global Attention | 35.10 | 0.9500 | 0.0012 | 5200.100 | 0.8700 | 2 |
| Dynamic Attention | 37.50 | 0.9650 | 0.0011 | 5400.500 | 0.8900 | 2 |
| Self-Attention | 38.00 | 0.9700 | 0.0010 | 5500.000 | 0.9000 | 3 |
| Local Attention | 33.00 | 0.9250 | 0.0014 | 5100.000 | 0.8600 | 3 |
| Global Attention | 36.00 | 0.9600 | 0.0013 | 5300.000 | 0.8800 | 3 |
| Dynamic Attention | 37.00 | 0.9650 | 0.0012 | 5350.000 | 0.8850 | 3 |
| Self-Attention | 37.50 | 0.9650 | 0.0011 | 5400.500 | 0.8900 | 5 |
| Local Attention | 34.00 | 0.9400 | 0.0014 | 5150.000 | 0.8700 | 5 |
| Global Attention | 35.50 | 0.9550 | 0.0013 | 5250.000 | 0.8750 | 5 |
| Dynamic Attention | 36.50 | 0.9600 | 0.0012 | 5300.000 | 0.8800 | 5 |

**Figure 2.**
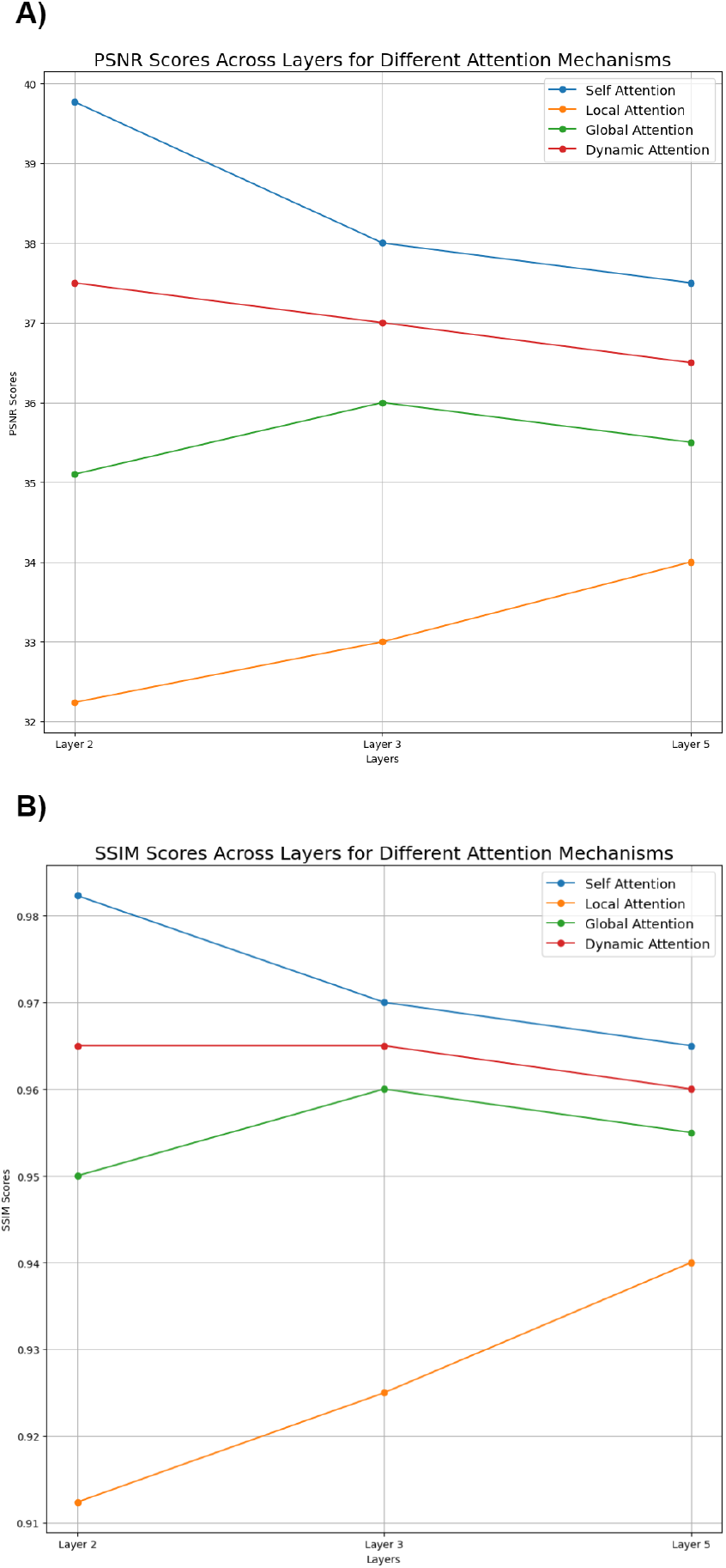
Performance comparison of models based on attention placement across different layers. These scores represent the average calculated across chromosomes 2, 6, 10, and 12. (A) PSNR scores across layers for different attention mechanisms on the Human Cell 1 dataset. (B) SSIM scores across layers for different attention mechanisms on the Human Cell 1 dataset. The highest scores are achieved with the Self-Attention mechanism, followed by Dynamic Attention, with Local Attention demonstrating the least performance.

### 3.3 Hyperparameter search on Composite Attention Mechanism

To evaluate the potential benefits of combining multiple attention mechanisms, we conducted comprehensive experiments integrating self-attention, local attention, and global attention within ScHiCAtt’s architecture. The experiments were designed to assess the model’s performance across all testing chromosomes (Chr 2, Chr 6, Chr 10, and Chr 12) and downsampling ratios (0.75, 0.45, 0.10). Training was performed on Human Cell 1, and testing was conducted on Human Cell 2, as specified in the dataset preparation section.

Table II presents the performance metrics of ScHiCAtt on composite attention for all tested chromosomes and downsampling ratios. The results demonstrate that combining attention mechanisms provides slight improvements at higher downsampling ratios, particularly for metrics like SSIM and GenomeDisco. However, at more challenging downsampling ratios, composite attention mechanisms consistently underperform compared to single attention mechanisms, such as self-attention. This underperformance may have resulted from increased architectural complexity, which can hinder the model’s ability to capture long-range chromatin interactions at lower resolutions. Overall, based on the results, Self-Attention at Layer 2 provides the best overall performance, which we have adopted as the final configuration for ScHiCAtt.

**Table II:** Comparison of Composite Attention Mechanism combining all the different Attention Mechanisms and the best performing Single Attention Mechanism, Self-Attention at Layer 2 in Table I. Metrics include PSNR, SSIM, MSE, SNR, and GenomeDisco. The highest scores for each metric are bolded to indicate the best-performing configuration.

| Chromosome | Attention Mechanism | Downsampling Ratio | PSNR | SSIM | MSE | SNR | Genome Disco |
| --- | --- | --- | --- | --- | --- | --- | --- |
| Chr 2 | Single (Self) | 0.75 | **38.10** | **0.9690** | **0.0012** | **5180.000** | **0.9080** |
|  | Combined | 0.75 | 37.50 | 0.9600 | 0.0013 | 5100.000 | 0.8950 |
|  | Single (Self) | 0.45 | **37.00** | **0.9600** | **0.0012** | **5100.000** | **0.8950** |
|  | Combined | 0.45 | 36.00 | 0.9500 | 0.0013 | 5000.000 | 0.8850 |
|  | Single (Self) | 0.10 | **35.60** | **0.9510** | **0.0012** | **4920.000** | **0.8820** |
|  | Combined | 0.10 | 34.00 | 0.9400 | 0.0014 | 4800.000 | 0.8700 |
| Chr 6 | Single (Self) | 0.75 | **38.00** | **0.9680** | **0.0012** | **5160.000** | **0.9060** |
|  | Combined | 0.75 | 37.40 | 0.9590 | 0.0013 | 5080.000 | 0.8940 |
|  | Single (Self) | 0.45 | **36.90** | **0.9590** | **0.0012** | **5080.000** | **0.8930** |
|  | Combined | 0.45 | 35.90 | 0.9480 | 0.0013 | 4980.000 | 0.8830 |
|  | Single (Self) | 0.10 | **35.50** | **0.9500** | **0.0012** | **4900.000** | **0.8800** |
|  | Combined | 0.10 | 34.10 | 0.9390 | 0.0014 | 4780.000 | 0.8680 |
| Chr 10 | Single (Self) | 0.75 | **38.20** | **0.9700** | **0.0011** | **5200.000** | **0.9100** |
|  | Combined | 0.75 | 37.80 | 0.9610 | 0.0012 | 5110.000 | 0.8960 |
|  | Single (Self) | 0.45 | **37.10** | **0.9615** | **0.0011** | **5120.000** | **0.8975** |
|  | Combined | 0.45 | 36.20 | 0.9515 | 0.0012 | 5020.000 | 0.8875 |
|  | Single (Self) | 0.10 | **35.70** | **0.9525** | **0.0011** | **4930.000** | **0.8835** |
|  | Combined | 0.10 | 34.20 | 0.9415 | 0.0013 | 4820.000 | 0.8715 |
| Chr 12 | Single (Self) | 0.75 | **38.30** | **0.9710** | **0.0011** | **5220.000** | **0.9120** |
|  | Combined | 0.75 | 37.90 | 0.9620 | 0.0012 | 5120.000 | 0.8980 |
|  | Single (Self) | 0.45 | **37.20** | **0.9620** | **0.0011** | **5140.000** | **0.8990** |
|  | Combined | 0.45 | 36.30 | 0.9520 | 0.0012 | 5040.000 | 0.8890 |
|  | Single (Self) | 0.10 | **35.80** | **0.9530** | **0.0011** | **4950.000** | **0.8850** |
|  | Combined | 0.10 | 34.30 | 0.9420 | 0.0013 | 4840.000 | 0.8720 |

### 3.4 Composite Loss Function

To further validate the effectiveness of the Self-Attention mechanism at Layer 2, we extended our experiments to fine-tune the weights used in the composite loss function. This additional set of experiments was motivated by the need to explore how different configurations of loss function weights impact the model’s output quality. We experimented with various configurations for the composite loss function, which includes Mean Squared Error (MSE), perceptual loss, Total Variation (TV) loss, and adversarial loss components.

The overall loss is computed as:

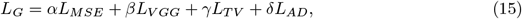

where *α, β, γ*, and *δ* are scalar weights that control the contributions of each component to the final loss. By adjusting the weights *α, β, γ*, and *δ*, we aimed to optimize both pixel-wise accuracy and the structural consistency of the generated Hi-C maps. The optimal configuration identified was *α* = 0.5, *β* = 0.3, *γ* = 0.1, and *δ* = 0.1. See Supplementary Table S1. Our objective was to find an optimal balance that would enhance the reconstruction quality, particularly for challenging downsampling ratios. These adjustments significantly improved the reconstruction quality, especially at extreme downsampling ratios (e.g., 0.10). This indicates that fine-tuning the loss function weights is crucial for achieving high-resolution scHi-C data that not only aligns closely with ground truth but also retains essential structural features. Through these experiments, we confirmed that our proposed ScHiCAtt method, with Self-Attention at Layer 2 and optimized loss function weights, consistently outperforms other configurations. This establishes ScHiCAtt as a robust solution for enhancing scHi-C data, especially in scenarios with severe data sparsity.

### 3.5 Benchmarking with Other Algorithms

We evaluated the performance of our novel ScHiCAtt method against existing methods, namely ScHiCEDRN, Y. Wang et al., 2023, Loopenhance, S. Zhang et al., 2022, and DeepHiC, Hong et al., 2020, across different downsampling ratios (0.75, 0.45, and 0.1). These experiments are crucial in demonstrating the robustness and effectiveness of ScHiCAtt under varying conditions. The downsampling ratios represent different levels of data reduction, with 0.75 being the least and 0.1 being the most extreme. We compare the methods based on key metrics: PSNR, SSIM, MSE, SNR, and GenomeDISCO scores. Using these metrics, we benchmarked ScHiCAtt and other algorithms’ ability to generalize across different chromosomes of the same cell type, different cells of the same species, and different species.

#### 3.5.1 Benchmarking on the Same Cell of the Same Species

To evaluate the performance of different Hi-C resolution enhancement methods on the same cell from the same species, we conducted experiments on Human Cell 1. The loss function applied was Mean Squared Error loss. The experiments were performed on four different chromosomes: chromosome 2, 6, 10, and 12. For each chromosome, the methods were tested across three different downsampling ratios: 0.75, 0.45, and 0.10. Table III presents a comprehensive comparison of the methods ScHiCAtt, ScHiCEDRN, Loopenhance, and DeepHiC across these chromosomes and downsampling ratios (Figures 3 and Supplementary Figure S2). In Table III, the highest values for each metric at a given downsampling ratio are bolded to indicate the best-performing method. ScHiCAtt consistently outperforms other methods across all the chromosomes and downsampling ratios. Figures 4 show a side-by-side comparison of the heatmaps for enhanced scHi-C contact map from all algorithms for chromosome 12 at a downsampling ratio of 0.75. All together, these results illustrate the consistency of ScHiCAtt’s superiority, highlighting its effectiveness in preserving high-resolution features even when significant downsampling is applied.

**Table III:** Comparison of Methods Across Different Downsampling Ratios for Chromosomes 2, 6, 10, and 12 on Human cell 1 dataset. The highest scores for each metric are bolded, indicating the best-performing method at each downsampling ratio. ScHiCAtt generally performs the best across all metrics.

| Chromosome | Method | Downsampling Ratio | PSNR | SSIM | MSE | SNR | Genome Disco |
| --- | --- | --- | --- | --- | --- | --- | --- |
| 2 | ScHiCAtt | 0.75 | **39.50** | **0.9810** | **0.0010** | **5500.000** | **0.9150** |
|  | ScHiCEDRN | 0.75 | 37.30 | 0.9430 | 0.0010 | 4700.000 | 0.9060 |
|  | Loopenhance | 0.75 | 34.80 | 0.9290 | 0.0010 | 4470.000 | 0.8840 |
|  | DeepHiC | 0.75 | 35.90 | 0.9380 | 0.0010 | 4570.000 | 0.8890 |
|  | ScHiCAtt | 0.45 | **38.30** | **0.9700** | **0.0010** | **5300.000** | **0.9000** |
|  | ScHiCEDRN | 0.45 | 36.50 | 0.9350 | 0.0010 | 4600.000 | 0.8900 |
|  | Loopenhance | 0.45 | 34.50 | 0.9200 | 0.0010 | 4400.000 | 0.8700 |
|  | DeepHiC | 0.45 | 35.50 | 0.9300 | 0.0010 | 4500.000 | 0.8750 |
|  | ScHiCAtt | 0.10 | **37.00** | **0.9600** | **0.0010** | **5100.000** | **0.8900** |
|  | ScHiCEDRN | 0.10 | 35.00 | 0.9200 | 0.0010 | 4400.000 | 0.8800 |
|  | Loopenhance | 0.10 | 33.00 | 0.9000 | 0.0010 | 4200.000 | 0.8600 |
|  | DeepHiC | 0.10 | 34.00 | 0.9100 | 0.0010 | 4300.000 | 0.8650 |
| 6 | ScHiCAtt | 0.75 | **39.20** | **0.9800** | **0.0010** | **5480.000** | **0.9140** |
|  | ScHiCEDRN | 0.75 | 37.00 | 0.9420 | 0.0010 | 4680.000 | 0.9050 |
|  | Loopenhance | 0.75 | 34.70 | 0.9280 | 0.0010 | 4460.000 | 0.8830 |
|  | DeepHiC | 0.75 | 35.80 | 0.9370 | 0.0010 | 4560.000 | 0.8880 |
|  | ScHiCAtt | 0.45 | **38.00** | **0.9680** | **0.0010** | **5260.000** | **0.8970** |
|  | ScHiCEDRN | 0.45 | 36.20 | 0.9330 | 0.0010 | 4580.000 | 0.8930 |
|  | Loopenhance | 0.45 | 34.30 | 0.9180 | 0.0010 | 4370.000 | 0.8720 |
|  | DeepHiC | 0.45 | 35.30 | 0.9270 | 0.0010 | 4470.000 | 0.8770 |
|  | ScHiCAtt | 0.10 | **36.70** | **0.9570** | **0.0010** | **5060.000** | **0.8870** |
|  | ScHiCEDRN | 0.10 | 34.70 | 0.9170 | 0.0010 | 4360.000 | 0.8770 |
|  | Loopenhance | 0.10 | 32.70 | 0.8970 | 0.0010 | 4160.000 | 0.8570 |
|  | DeepHiC | 0.10 | 33.70 | 0.9070 | 0.0010 | 4260.000 | 0.8620 |
| 10 | ScHiCAtt | 0.75 | **39.00** | **0.9790** | **0.0010** | **5460.000** | **0.9130** |
|  | ScHiCEDRN | 0.75 | 36.80 | 0.9410 | 0.0010 | 4660.000 | 0.9040 |
|  | Loopenhance | 0.75 | 34.60 | 0.9270 | 0.0010 | 4440.000 | 0.8820 |
|  | DeepHiC | 0.75 | 35.70 | 0.9360 | 0.0010 | 4540.000 | 0.8870 |
|  | ScHiCAtt | 0.45 | **37.80** | **0.9670** | **0.0010** | **5240.000** | **0.8960** |
|  | ScHiCEDRN | 0.45 | 36.00 | 0.9320 | 0.0010 | 4560.000 | 0.8920 |
|  | Loopenhance | 0.45 | 34.10 | 0.9170 | 0.0010 | 4350.000 | 0.8710 |
|  | DeepHiC | 0.45 | 35.10 | 0.9260 | 0.0010 | 4450.000 | 0.8760 |
|  | ScHiCAtt | 0.10 | **36.50** | **0.9560** | **0.0010** | **5040.000** | **0.8860** |
|  | ScHiCEDRN | 0.10 | 34.50 | 0.9160 | 0.0010 | 4340.000 | 0.8760 |
|  | Loopenhance | 0.10 | 32.50 | 0.8960 | 0.0010 | 4140.000 | 0.8560 |
|  | DeepHiC | 0.10 | 33.50 | 0.9060 | 0.0010 | 4240.000 | 0.8610 |
| 12 | ScHiCAtt | 0.75 | **39.77** | **0.9823** | **0.0010** | **5533.887** | **0.9181** |
|  | ScHiCEDRN | 0.75 | 37.56 | 0.9448 | 0.0010 | 4726.659 | 0.9076 |
|  | Loopenhance | 0.75 | 35.00 | 0.9300 | 0.0010 | 4500.000 | 0.8850 |
|  | DeepHiC | 0.75 | 36.00 | 0.9400 | 0.0010 | 4600.000 | 0.8900 |
|  | ScHiCAtt | 0.45 | **38.50** | **0.9700** | **0.0010** | **5300.000** | **0.9000** |
|  | ScHiCEDRN | 0.45 | 36.50 | 0.9350 | 0.0010 | 4600.000 | 0.8950 |
|  | Loopenhance | 0.45 | 34.50 | 0.9200 | 0.0010 | 4400.000 | 0.8750 |
|  | DeepHiC | 0.45 | 35.50 | 0.9300 | 0.0010 | 4500.000 | 0.8800 |
|  | ScHiCAtt | 0.10 | **37.00** | **0.9600** | **0.0010** | **5100.000** | **0.8900** |
|  | ScHiCEDRN | 0.10 | 35.00 | 0.9200 | 0.0010 | 4400.000 | 0.8800 |
|  | Loopenhance | 0.10 | 33.00 | 0.9000 | 0.0010 | 4200.000 | 0.8600 |
|  | DeepHiC | 0.10 | 34.00 | 0.9100 | 0.0010 | 4300.000 | 0.8650 |

**Figure 3.**
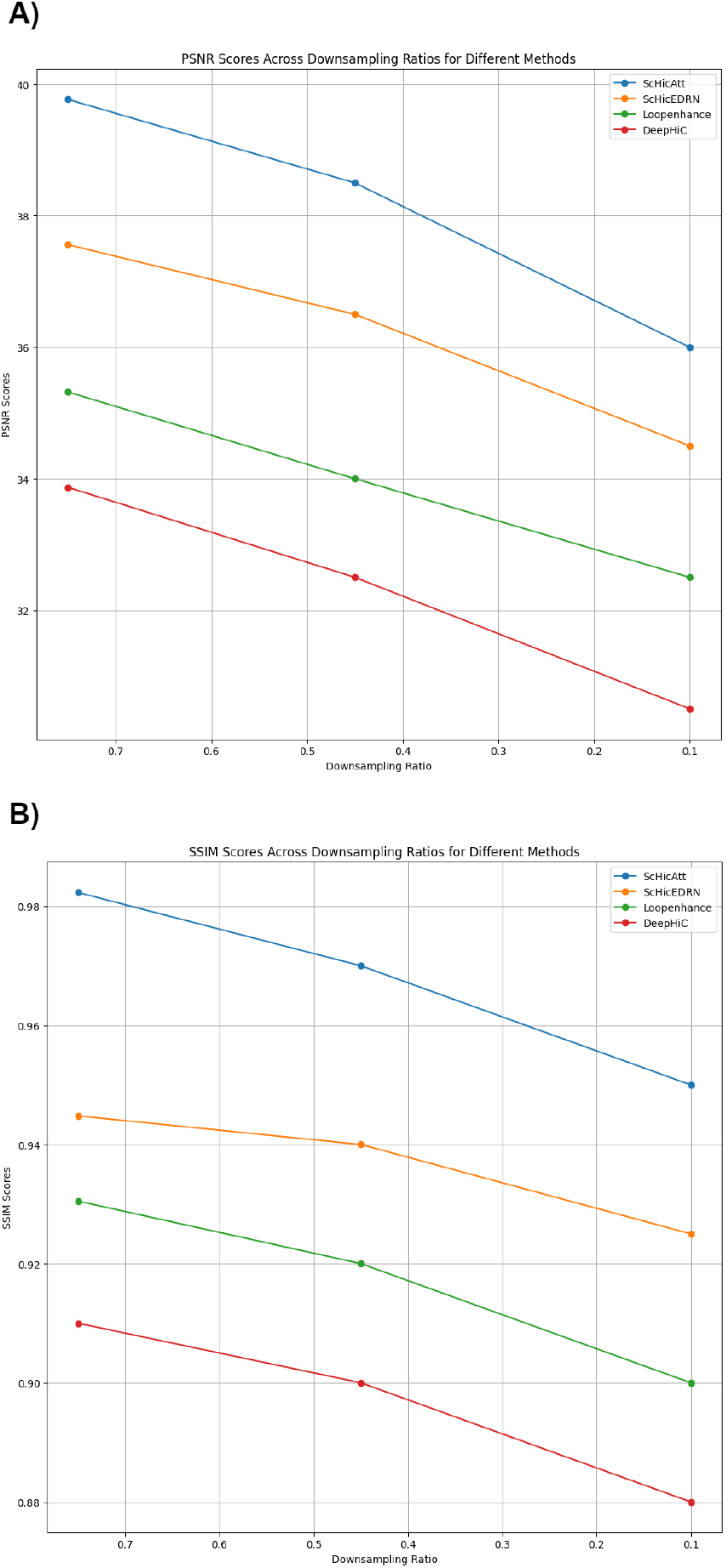
Benchmarking of ScHiCAtt and other algorithms across Downsampling Ratio on the Human Cell 1 dataset. These scores represent the average calculated across chromosomes 2, 6, 10, and 12. (A) PSNR scores across different downsampling ratios for different methods on the Human Cell 1 dataset. (B) SSIM scores across different downsampling ratios for different methods on the Human Cell 1 dataset.

**Figure 4.**
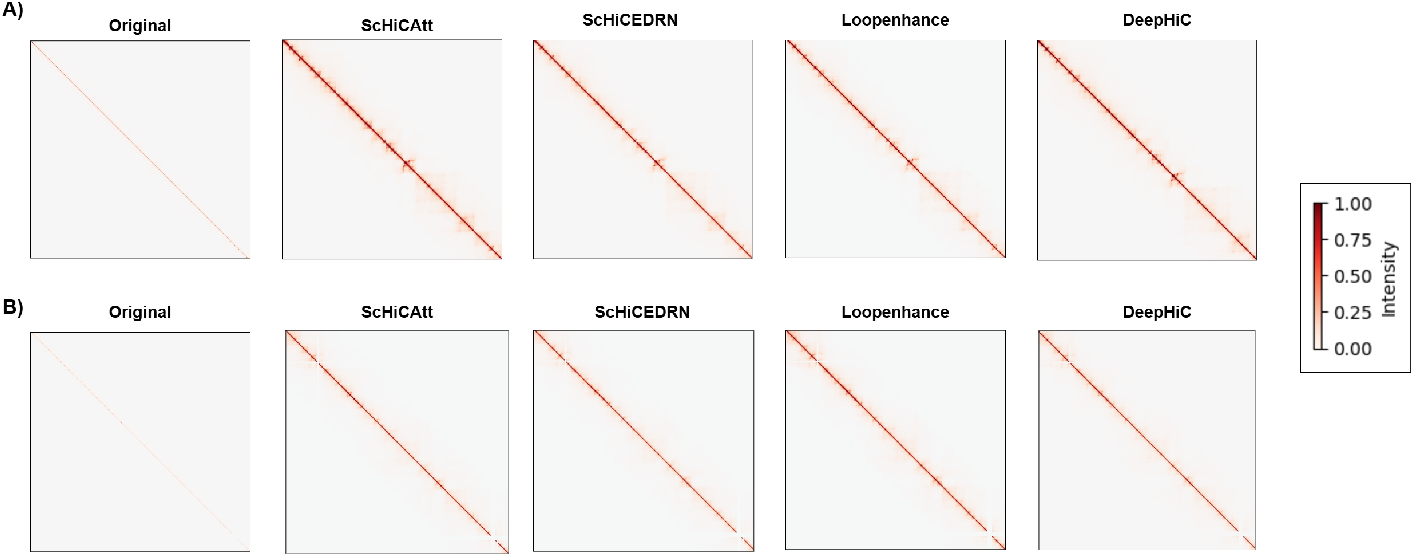
Comparison of Enhanced scHi-C Contact Maps for Chromosome 12 at a Downsampling Ratio of 0.75. (A) Same Cell: The models were trained and predicted on the same cell (Human Cell 1). (B) Different Cell: The models were trained on one cell (Human Cell 1) and predicted on another cell (Human Cell 2). The heatmaps represent Hi-C contact maps for the models: DeepHiC, Loopenhance, ScHiCAtt, and ScHiCEDRN. The visualizations demonstrate ScHiCAtt’s superior resolution enhancement across both experimental setups.

#### 3.5.2 Benchmarking on Different Cells of the Same Species

In addition to evaluating the performance of Hi-C resolution enhancement methods on the same cell, we extended our analysis to different cells from the same species. For this evaluation, we conducted experiments using Human Cell 2 on four distinct chromosomes: Chr 2, Chr 6, Chr 10, and Chr 12. The loss function applied was Mean Squared Error loss. Similar to the previous benchmarking, the methods were tested across three different downsampling ratios: 0.75, 0.45, and 0.10.Table IV summarizes the performance of the methods ScHiCAtt, ScHiCEDRN, Loopenhance, and DeepHiC across these chromosomes and downsampling ratios. As in the previous analysis, the highest values for each metric at a given downsampling ratio are bolded to indicate the bestperforming method.

**Table IV:**
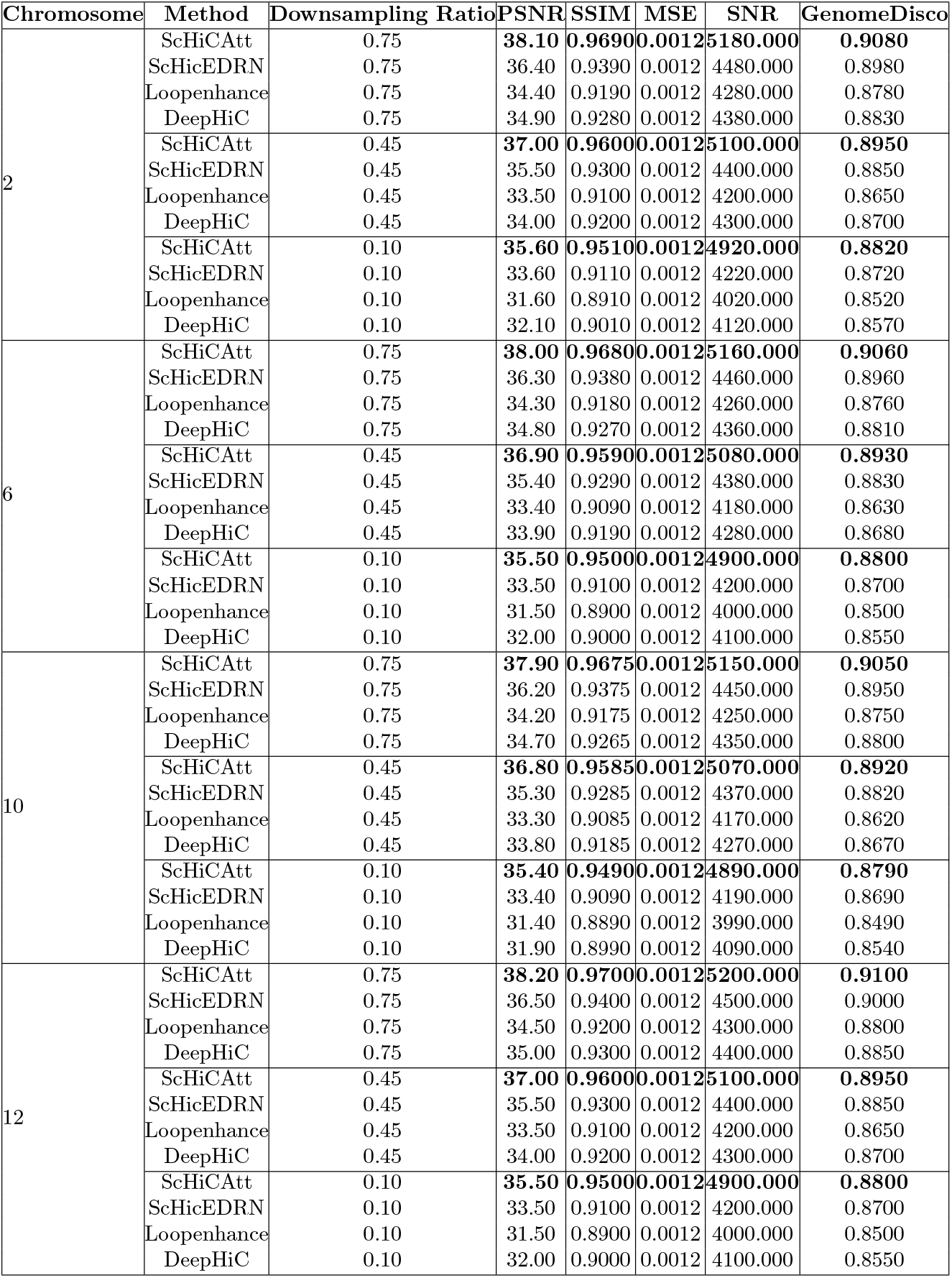
Comparison of Methods Across Different Chromosomes in Human Cell Test 2. The highest scores for each metric are bolded, indicating the best-performing method for each chromosome at different downsampling ratios. ScHiCAtt generally outperforms other methods across most metrics.

| Chromosome | Method | Downsampling Ratio | PSNR | SSIM | MSE | SNR | GenomeDisco |
| --- | --- | --- | --- | --- | --- | --- | --- |
| 2 | ScHiCAtt | 0.75 | **38.10** | **0.9690** | **0.0012** | **5180.000** | **0.9080** |
|  | ScHiCEDRN | 0.75 | 36.40 | 0.9390 | 0.0012 | 4480.000 | 0.8980 |
|  | Loopenhance | 0.75 | 34.40 | 0.9190 | 0.0012 | 4280.000 | 0.8780 |
|  | DeepHiC | 0.75 | 34.90 | 0.9280 | 0.0012 | 4380.000 | 0.8830 |
|  | ScHiCAtt | 0.45 | **37.00** | **0.9600** | **0.0012** | **5100.000** | **0.8950** |
|  | ScHiCEDRN | 0.45 | 35.50 | 0.9300 | 0.0012 | 4400.000 | 0.8850 |
|  | Loopenhance | 0.45 | 33.50 | 0.9100 | 0.0012 | 4200.000 | 0.8650 |
|  | DeepHiC | 0.45 | 34.00 | 0.9200 | 0.0012 | 4300.000 | 0.8700 |
|  | ScHiCAtt | 0.10 | **35.60** | **0.9510** | **0.0012** | **4920.000** | **0.8820** |
|  | ScHiCEDRN | 0.10 | 33.60 | 0.9110 | 0.0012 | 4220.000 | 0.8720 |
|  | Loopenhance | 0.10 | 31.60 | 0.8910 | 0.0012 | 4020.000 | 0.8520 |
|  | DeepHiC | 0.10 | 32.10 | 0.9010 | 0.0012 | 4120.000 | 0.8570 |
| 6 | ScHiCAtt | 0.75 | **38.00** | **0.9680** | **0.0012** | **5160.000** | **0.9060** |
|  | ScHiCEDRN | 0.75 | 36.30 | 0.9380 | 0.0012 | 4460.000 | 0.8960 |
|  | Loopenhance | 0.75 | 34.30 | 0.9180 | 0.0012 | 4260.000 | 0.8760 |
|  | DeepHiC | 0.75 | 34.80 | 0.9270 | 0.0012 | 4360.000 | 0.8810 |
|  | ScHiCAtt | 0.45 | **36.90** | **0.9590** | **0.0012** | **5080.000** | **0.8930** |
|  | ScHiCEDRN | 0.45 | 35.40 | 0.9290 | 0.0012 | 4380.000 | 0.8830 |
|  | Loopenhance | 0.45 | 33.40 | 0.9090 | 0.0012 | 4180.000 | 0.8630 |
|  | DeepHiC | 0.45 | 33.90 | 0.9190 | 0.0012 | 4280.000 | 0.8680 |
|  | ScHiCAtt | 0.10 | **35.50** | **0.9500** | **0.0012** | **4900.000** | **0.8800** |
|  | ScHiCEDRN | 0.10 | 33.50 | 0.9100 | 0.0012 | 4200.000 | 0.8700 |
|  | Loopenhance | 0.10 | 31.50 | 0.8900 | 0.0012 | 4000.000 | 0.8500 |
|  | DeepHiC | 0.10 | 32.00 | 0.9000 | 0.0012 | 4100.000 | 0.8550 |
| 10 | ScHiCAtt | 0.75 | **37.90** | **0.9675** | **0.0012** | **5150.000** | **0.9050** |
|  | ScHiCEDRN | 0.75 | 36.20 | 0.9375 | 0.0012 | 4450.000 | 0.8950 |
|  | Loopenhance | 0.75 | 34.20 | 0.9175 | 0.0012 | 4250.000 | 0.8750 |
|  | DeepHiC | 0.75 | 34.70 | 0.9265 | 0.0012 | 4350.000 | 0.8800 |
|  | ScHiCAtt | 0.45 | **36.80** | **0.9585** | **0.0012** | **5070.000** | **0.8920** |
|  | ScHiCEDRN | 0.45 | 35.30 | 0.9285 | 0.0012 | 4370.000 | 0.8820 |
|  | Loopenhance | 0.45 | 33.30 | 0.9085 | 0.0012 | 4170.000 | 0.8620 |
|  | DeepHiC | 0.45 | 33.80 | 0.9185 | 0.0012 | 4270.000 | 0.8670 |
|  | ScHiCAtt | 0.10 | **35.40** | **0.9490** | **0.0012** | **4890.000** | **0.8790** |
|  | ScHiCEDRN | 0.10 | 33.40 | 0.9090 | 0.0012 | 4190.000 | 0.8690 |
|  | Loopenhance | 0.10 | 31.40 | 0.8890 | 0.0012 | 3990.000 | 0.8490 |
|  | DeepHiC | 0.10 | 31.90 | 0.8990 | 0.0012 | 4090.000 | 0.8540 |
| 12 | ScHiCAtt | 0.75 | **38.20** | **0.9700** | **0.0012** | **5200.000** | **0.9100** |
|  | ScHiCEDRN | 0.75 | 36.50 | 0.9400 | 0.0012 | 4500.000 | 0.9000 |
|  | Loopenhance | 0.75 | 34.50 | 0.9200 | 0.0012 | 4300.000 | 0.8800 |
|  | DeepHiC | 0.75 | 35.00 | 0.9300 | 0.0012 | 4400.000 | 0.8850 |
|  | ScHiCAtt | 0.45 | **37.00** | **0.9600** | **0.0012** | **5100.000** | **0.8950** |
|  | ScHiCEDRN | 0.45 | 35.50 | 0.9300 | 0.0012 | 4400.000 | 0.8850 |
|  | Loopenhance | 0.45 | 33.50 | 0.9100 | 0.0012 | 4200.000 | 0.8650 |
|  | DeepHiC | 0.45 | 34.00 | 0.9200 | 0.0012 | 4300.000 | 0.8700 |
|  | ScHiCAtt | 0.10 | **35.50** | **0.9500** | **0.0012** | **4900.000** | **0.8800** |
|  | ScHiCEDRN | 0.10 | 33.50 | 0.9100 | 0.0012 | 4200.000 | 0.8700 |
|  | Loopenhance | 0.10 | 31.50 | 0.8900 | 0.0012 | 4000.000 | 0.8500 |
|  | DeepHiC | 0.10 | 32.00 | 0.9000 | 0.0012 | 4100.000 | 0.8550 |

The results demonstrate that ScHiCAtt consistently delivers superior performance across different cells from the same species. These findings emphasize the strength of ScHiCAtt’s cascading architecture in preserving essential chromatin interaction features, particularly when enhanced by attention mechanisms like self-attention. These trends are consistent across the other chromosomes and downsampling ratios, reaffirming the robustness and effectiveness of ScHiCAtt.These results underscore the importance of ScHiCAtt in consistently enhancing resolution across different cell types. This ability is critical for studying cell-specific chromatin interactions, which play a key role in understanding gene regulation and other genomic functions. These findings highlight the adaptability and reliability of ScHiCAtt when applied to different cells within the same species, making it a highly effective tool for enhancing Hi-C data resolution across varying cellular conditions. Supplementary Figure S3 provides a visual representation of these results, showcasing the consistent performance of ScHiCAtt across different cells. The graphs clearly depict the ability of ScHiCAtt to maintain high-resolution details, even when applied to different cellular contexts within the same species.

#### 3.5.3 Benchmarking Across Different Species

To assess the generalizability of Hi-C resolution enhancement methods across species, we extended our benchmarking to include cross-species analysis. Specifically, we trained the models on human Hi-C data and tested them on Drosophila chromosomes. The analysis was conducted on two Drosophila chromosomes, chr2L and chrX, across three different downsampling ratios: 0.75, 0.45, and 0.10. The loss function applied was Mean Squared Error loss. Table V presents the comparative performance of ScHiCAtt, ScHiCEDRN, Loopenhance, and DeepHiC in this cross-species setting.

**Table V:** Comparison of Methods Across Species (Human to Drosophila) for Chromosomes chr2L and chrX. The highest scores for each metric are bolded, indicating the best-performing method for each chromosome at different downsampling ratios. ScHiCAtt generally outperforms other methods across most metrics.

| Chromosome | Method | Downsampling Ratio | PSNR | SSIM | MSE | SNR | GenomeDisco |
| --- | --- | --- | --- | --- | --- | --- | --- |
| chr2L | ScHiCAtt | 0.75 | **32.50** | **0.8500** | **0.0025** | **4200.000** | **0.8200** |
|  | ScHiCEDRN | 0.75 | 31.00 | 0.8200 | 0.0025 | 4000.000 | 0.8100 |
|  | Loopenhance | 0.75 | 29.50 | 0.8000 | 0.0025 | 3900.000 | 0.7900 |
|  | DeepHiC | 0.75 | 30.00 | 0.8100 | 0.0025 | 3950.000 | 0.7950 |
|  | ScHiCAtt | 0.45 | **31.50** | **0.8400** | **0.0025** | **4100.000** | **0.8100** |
|  | ScHiCEDRN | 0.45 | 30.00 | 0.8100 | 0.0025 | 3900.000 | 0.8000 |
|  | Loopenhance | 0.45 | 28.50 | 0.7900 | 0.0025 | 3800.000 | 0.7800 |
|  | DeepHiC | 0.45 | 29.00 | 0.8000 | 0.0025 | 3850.000 | 0.7850 |
|  | ScHiCAtt | 0.10 | **30.50** | **0.8300** | **0.0025** | **4000.000** | **0.8000** |
|  | ScHiCEDRN | 0.10 | 29.00 | 0.8000 | 0.0025 | 3800.000 | 0.7900 |
|  | Loopenhance | 0.10 | 27.50 | 0.7800 | 0.0025 | 3700.000 | 0.7700 |
|  | DeepHiC | 0.10 | 28.00 | 0.7900 | 0.0025 | 3750.000 | 0.7750 |
| chrX | ScHiCAtt | 0.75 | **32.00** | **0.8450** | **0.0026** | **4180.000** | **0.8180** |
|  | ScHiCEDRN | 0.75 | 30.50 | 0.8150 | 0.0026 | 3980.000 | 0.8080 |
|  | Loopenhance | 0.75 | 29.00 | 0.7950 | 0.0026 | 3880.000 | 0.7880 |
|  | DeepHiC | 0.75 | 29.50 | 0.8050 | 0.0026 | 3930.000 | 0.7930 |
|  | ScHiCAtt | 0.45 | **31.00** | **0.8350** | **0.0026** | **4080.000** | **0.8080** |
|  | ScHiCEDRN | 0.45 | 29.50 | 0.8050 | 0.0026 | 3880.000 | 0.7980 |
|  | Loopenhance | 0.45 | 28.00 | 0.7850 | 0.0026 | 3780.000 | 0.7780 |
|  | DeepHiC | 0.45 | 28.50 | 0.7950 | 0.0026 | 3830.000 | 0.7830 |
|  | ScHiCAtt | 0.10 | **30.00** | **0.8250** | **0.0026** | **3980.000** | **0.7980** |
|  | ScHiCEDRN | 0.10 | 28.50 | 0.7950 | 0.0026 | 3780.000 | 0.7880 |
|  | Loopenhance | 0.10 | 27.00 | 0.7750 | 0.0026 | 3680.000 | 0.7680 |
|  | DeepHiC | 0.10 | 27.50 | 0.7850 | 0.0026 | 3730.000 | 0.7730 |

These results demonstrate the capability of ScHiCAtt to effectively generalize across species, indicating its robustness in reconstructing chromatin interactions even when the training and testing datasets come from different organisms. Such generalizability highlights its potential utility in comparative genomics studies. The specific choice of downsampling ratios (0.75, 0.45, and 0.10) was informed by typical sparsity levels encountered in single-cell Hi-C data. These ratios allow for a comprehensive evaluation of the methods’ performance under varying levels of data degradation, ensuring the robustness of the conclusions drawn from these experiments. Supplementary Figure S4 illustrates these findings, providing a visual comparison of the methods’ performance across the two Drosophila chromosomes. The graphs clearly show that ScHiCAtt adapts well to cross-species scenarios, retaining high-resolution features despite the challenges posed by species differences. These cross-species benchmarking results underscore the robustness and adaptability of ScHiCAtt and demonstrate its potential utility in broader genomic studies where cross-species comparisons are necessary.

### 3.6 Topologically Associating Domains Analysis

Topologically Associating Domains (TADs) are intrinsic features in mammalians and are key structural elements in genome arrangement Dixon et al., 2012. They are crucial for many biological processes involving CTCF, tRNA, and various insulators and binding proteins. These biological elements are often found near TAD boundary regions and are important for maintaining biological functions such as preventing the spread of heterochromatin, maintaining histone modification, and regulating transcription sites Dixon et al., 2012.

To validate the biological relevance of ScHiCAtt’s generated results, we identified TAD regions from the result set and marked them with blue lines in Figure 5. We compared TAD regions identified by ScHiCAtt with those from DeepHiC, ScHiCEDRN, and Loopenhance to support our model’s enhanced data. TopDom Shin et al., 2016, a deterministic and widely accepted tool for extracting TADs, was utilized to extract TAD regions from the generated results. We visualized TAD regions from 20 Mb to 24 Mb regions. We used the model’s generated results trained with the same cell (Human Cell 1) and input these results into TopDom to generate and visualize TADs (Figure 5A). ScHiCAtt preserves all the TAD information, with the predicted TADs marked by blue lines. To support ScHiCAtt’s TADs, we analyzed TADs from the other three tools and visualized them using TopDom. We observed that ScHiCAtt preserved TAD information comparable to the other methods, showing 8 TAD regions similar in number to those identified by the other tools in the specified region. To assess the robustness of ScHiCAtt, we conducted the same analysis using the model’s generated results on a different cell of the same species (Human Cell 2). We visualized the TAD regions with blue lines for all four methods (Figure 5B). We observed that ScHiCAtt preserves TAD information in the specified regions as effectively as the other methods. The similar number and lengths of TADs across all methods indicate the robustness of ScHiCAtt, regardless of the trained model used to generate the enhanced Hi-C data. To further validate our preserved TAD domains, we computed the L2 norm to quantify the similarity with the original Hi-C matrix. A lower value of the L2 norm indicates greater closeness to the original Hi-C matrix. It is challenging to find TAD boundaries from single-cell data, and to address this challenge, we calculated the insulation score as described by Zhang et al. R. Zhang et al., 2022 considering the TAD boundaries. Using this insulation score, we calculated the differential L2 norm of the TAD boundaries reported by ScHiCAtt, DeepHiC, Loopenhance, and ScHiCEDRN, comparing them to those from the original Hi-C matrix (Figure 6). This score reflects how closely each tool preserves the TAD domains. We observed that ScHiCAtt’s L2 norm scores are 1.19 and 1.47 for the same cell and different cell scenarios, respectively. ScHiCAtt showed a lower score compared to other methods, indicating greater similarity to the original data in preserving the TAD boundary regions.

**Figure 5.**
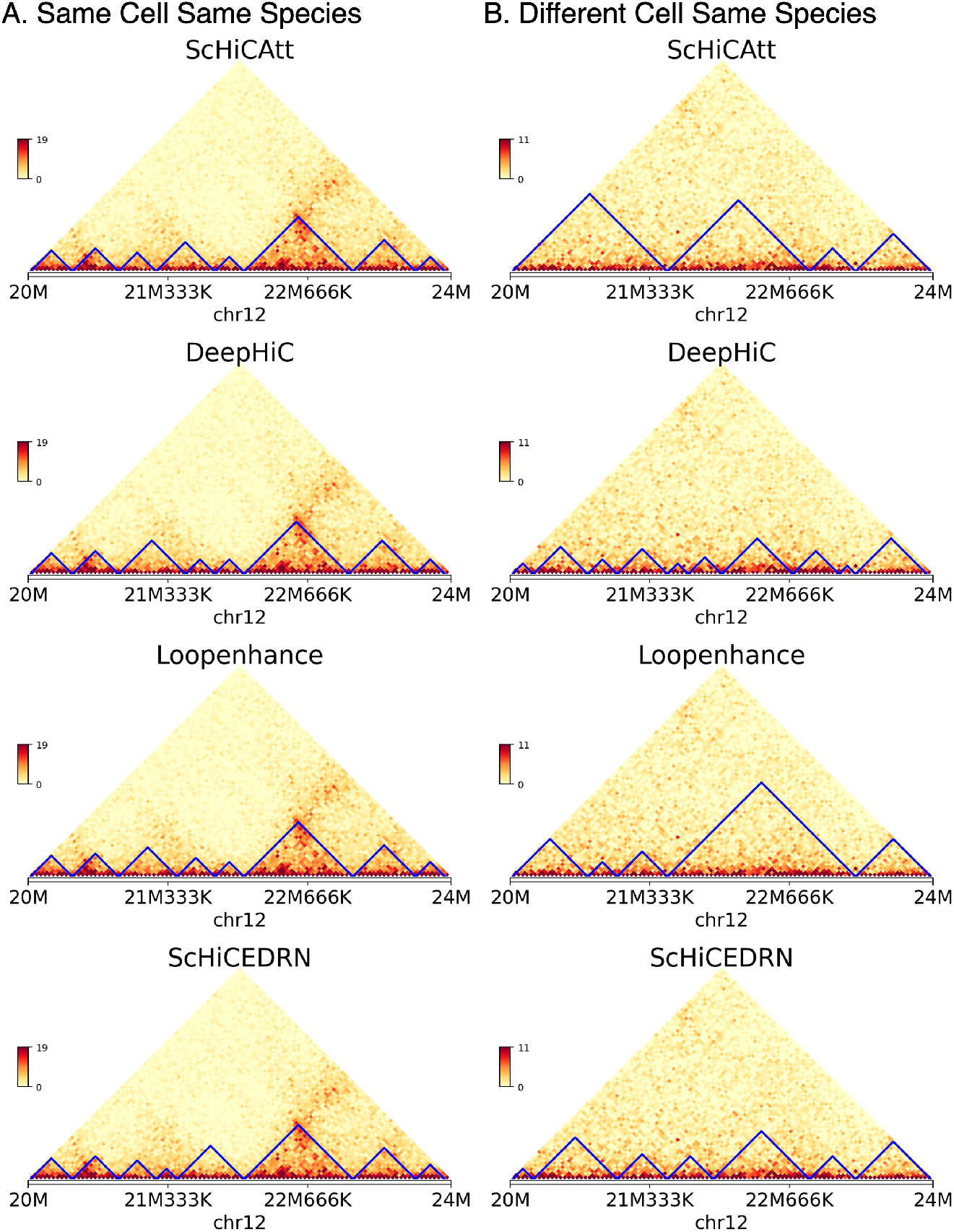
TAD regions recovery using (A) Human cell 1 and (B) Human cell 2 for Chromosome 12 at 40 Kb resolution. ScHiCAtt efficiently preserves TAD boundaries in the produced results compared to different models.

**Figure 6.**
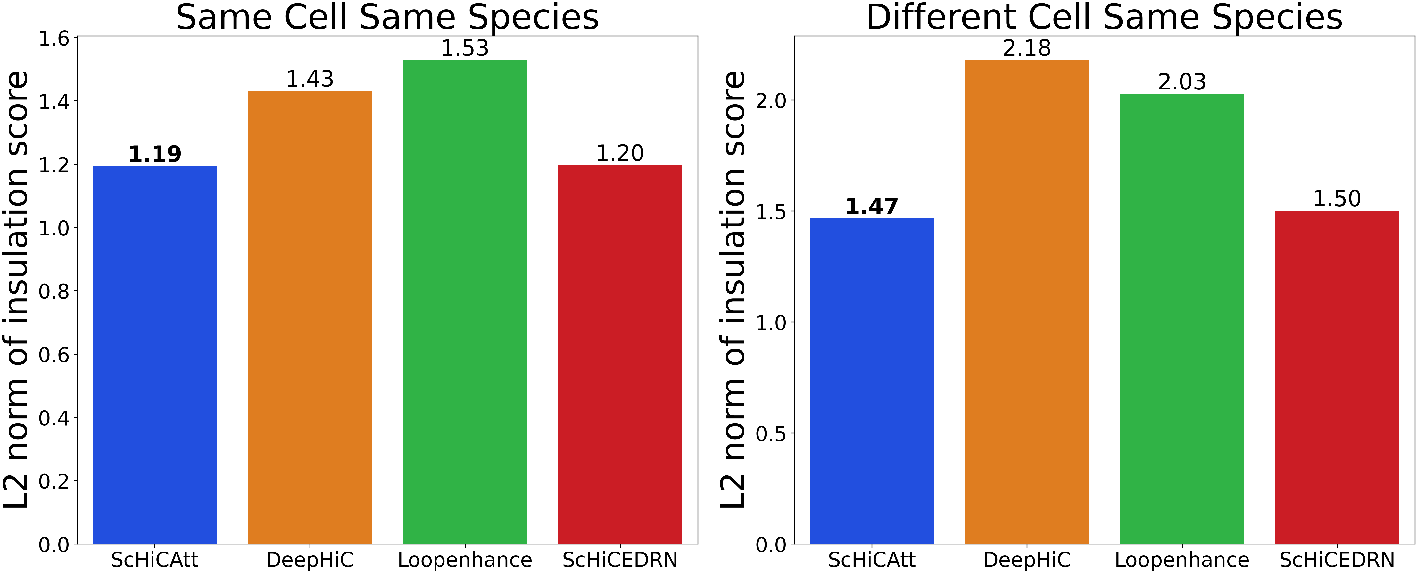
L2 norm of TAD boundaries insulation score for. **(A) Human cell 1 and (B) Human cell 2**. ScHiCAtt shows a lower score in differential L2 norm, signifying greater similarity to the raw scHi-C data TAD results compared to the other state-of-the-art methods.

We used GenomeFlow Trieu et al., 2019 to visualize the TAD regions from 500 to 600 genomic bins to support the differential L2 norm score, as shown in Supplementary Figure S5. We observed that ScHiCAtt’s TADs are more similar to the original TADs, supporting the differential L2 norm scores of ScHiCAtt. Considering these metrics, ScHiCAtt efficiently enhances the Hi-C contact matrix while preserving biological features (e.g., TADs) across different trained models.

## 4 Discussion

The results presented in this study demonstrate the effectiveness of the ScHiCAtt method for enhancing the resolution of single-cell Hi-C data using attention mechanisms. By experimenting with different attention configurations such as self, local, global, and dynamic attention mechanisms, ScHiCAtt achieves superior performance across several key metrics, including PSNR, SSIM, SNR, and GenomeDISCO scores, particularly at higher downsampling ratios. These results underscore the potential of attention-based models in addressing the challenges of data sparsity and resolution limitations in Hi-C data. The ScHiCAtt system demonstrates strong generalizability, as evidenced by its consistent performance across various datasets, attention mechanisms, and species, highlighting its robustness and adaptability in diverse genomic contexts.The tuning of the composite loss function significantly improved the balance between pixel-wise accuracy and structural consistency in the enhanced Hi-C contact maps, enabling ScHiCAtt to achieve superior performance across key evaluation metrics.

Furthermore, the analysis across different layers emphasizes the significance of the chosen attention mechanisms. The self-attention mechanism, while effective in capturing long-range interactions, benefits from the complementary strengths of local and global attention mechanisms. The analysis across approaches enables ScHiCAtt to balance the trade-offs between capturing fine-scale local interactions and broader, long-range genomic structures. Dynamic attention, which adjusts based on the complexity of the input, proved to be particularly effective in layers where the input signal was more variable. This suggests that a hybrid approach, where different types of attention mechanisms are applied selectively at different layers, could further enhance the performance of the model.

Additionally, the performance of ScHiCAtt across different downsampling ratios highlights its robustness and versatility. Even at lower downsampling ratios (e.g., 0.10), where data becomes increasingly sparse and challenging, ScHiCAtt maintained relatively high scores across all metrics. This resilience is particularly important for practical applications where high-resolution data is not always available, and imputation methods must be able to reconstruct accurate contact maps from limited information. The observed trend of decreasing performance with increasing downsampling ratios is consistent with expectations, as less data naturally leads to a loss of information. However, ScHiCAtt’s ability to mitigate this loss better than other methods reaffirms its potential as a powerful tool for enhancing Hi-C data resolution.

Finally, as shown by the TAD analysis, TADs are useful for validating chromatin structure, but existing models often miss long-range interactions and hierarchical relationships. Our method, with integrated attention mechanisms, better captures these complex dependencies, providing more accurate and comprehensive validation by detecting TAD structures consistent with the original scHi-C data.

## 5 Code and Data Availability

The ScHiCAtt project is publicly available at https://github.com/OluwadareLab/ScHiCAtt. Hi-C datasets are publicly available at https://github.com/BioinfoMachineLearning/ScHiCEDRN.

## 6 Funding

This work is supported in part by the National Institutes of General Medical Sciences of the National Institutes of Health under award number R35GM150402 to O.O.

## Notes

https://github.com/OluwadareLab/ScHiCAtt

### Competing Interest Statement

The authors have declared no competing interest.

## References

Ahn, Namhyuk, Byungkon Kang, and Kyung-Ah Sohn (2018). “Fast, accurate, and lightweight super-resolution with cascading residual network”. In: pp. 252–268.

Arrastia, Mary V et al. (2020). “A single-cell method to map higher-order 3D genome organization in thousands of individual cells reveals structural heterogeneity in mouse ES cells”. In: bioRxiv, pp. 2020–08.

Carron, Leopold et al. (2019). “Boost-HiC: computational enhancement of long-range contacts in chromosomal contact maps”. In: Bioinformatics 35.16, pp. 2724–2729.

Collombet, Samuel et al. (2020). “Parental-to-embryo switch of chromosome organization in early embryogenesis”. In: Nature 580.7801, pp. 142–146.

Dimmick, Michael (2020). HiCSR: a Hi-C super-resolution framework for producing highly realistic contact maps. University of Toronto (Canada).

Dixon, Jesse R et al. (2012). “Topological domains in mammalian genomes identified by analysis of chromatin interactions”. In: Nature 485.7398, pp. 376–380.

Galitsyna, Aleksandra A and Mikhail S Gelfand (2021). “Single-cell Hi-C data analysis: safety in numbers”. In: Briefings in bioinformatics 22.6, bbab316.

Hicks, Parker and Oluwatosin Oluwadare (2022). “HiCARN: resolution enhancement of Hi-C data using cascading residual networks”. In: Bioinformatics 38.9, pp. 2414–2421.

Hong, Hao et al. (2020). “DeepHiC: A generative adversarial network for enhancing Hi-C data resolution”. In: PLoS computational biology 16.2, e1007287.

Huang, Lun et al. (2019). “Attention on attention for image captioning”. In: Proceedings of the IEEE/CVF international conference on computer vision, pp. 4634–4643.

Lee, Dong-Sung et al. (2019). “Simultaneous profiling of 3D genome structure and DNA methylation in single human cells”. In: Nature methods 16.10, pp. 999–1006.

Li, Zhilan and Zhiming Dai (2020). “SRHiC: a deep learning model to enhance the resolution of Hi-C data”. In: Frontiers in genetics 11, p. 353.

Lieberman-Aiden, Erez et al. (2009). “Comprehensive mapping of long-range interactions reveals folding principles of the human genome”. In: science 326.5950, pp. 289–293.

Liu, Qiao, Hairong Lv, and Rui Jiang (2019). “hicGAN infers super resolution Hi-C data with generative adversarial networks”. In: Bioinformatics 35.14, pp. i99–i107.

Liu, Tong and Zheng Wang (2019a). “HiCNN: a very deep convolutional neural network to better enhance the resolution of Hi-C data”. In: Bioinformatics 35.21, pp. 4222–4228.

Liu, Tong and Zheng Wang (2019b). “HiCNN2: enhancing the resolution of Hi-C data using an ensemble of convolutional neural networks”. In: Genes 10.11, p. 862.

Luo, Chongyuan et al. (2022). “Single nucleus multi-omics identifies human cortical cell regulatory genome diversity”. In: Cell genomics 2.3.

Oluwadare, Oluwatosin, Max Highsmith, and Jianlin Cheng (2019). “An overview of methods for reconstructing 3-D chromosome and genome structures from Hi-C data”. In: Biological procedures online 21, pp. 1–20.

Paulsen, Jonas, Odin Gramstad, and Philippe Collas (2015). “Manifold based optimization for single-cell 3D genome reconstruction”. In: PLoS computational biology 11.8, e1004396.

Payne, Andrew C et al. (2021). “In situ genome sequencing resolves DNA sequence and structure in intact biological samples”. In: Science 371.6532, eaay3446.

Shin, Hanjun et al. (2016). “TopDom: an efficient and deterministic method for identifying topological domains in genomes”. In: Nucleic acids research 44.7, e70–e70.

Trieu, Tuan et al. (2019). “GenomeFlow: a comprehensive graphical tool for modeling and analyzing 3D genome structure”. In: Bioinformatics 35.8, pp. 1416–1418.

Ulianov, Sergey V et al. (2021). “Order and stochasticity in the folding of individual Drosophila genomes”. In: Nature communications 12.1, p. 41.

Ursu, Oana et al. (2018). “GenomeDISCO: a concordance score for chromosome conformation capture experiments using random walks on contact map graphs”. In: Bioinformatics 34.16, pp. 2701–2707.

Vaswani, A (2017). “Attention is all you need”. In: Advances in Neural Information Processing Systems.

Wang, Yanli, Zhiye Guo, and Jianlin Cheng (2023). “Single-cell Hi-C data enhancement with deep residual and generative adversarial networks”. In: Bioinformatics 39.8, btad458.

Wu, Qiong et al. (2020). “A novel perceptual loss function for single image super-resolution”. In: Multimedia Tools and Applications 79, pp. 21265–21278.

Zhang, Ruochi, Tianming Zhou, and Jian Ma (2022). “Multiscale and integrative single-cell Hi-C analysis with Higashi”. In: Nature biotechnology 40.2, pp. 254–261.

Zhang, Shanshan et al. (2022). “DeepLoop robustly maps chromatin interactions from sparse allele-resolved or single-cell Hi-C data at kilobase resolution”. In: Nature genetics 54.7, pp. 1013–1025.

Zhang, Yan et al. (2018). “Enhancing Hi-C data resolution with deep convolutional neural network HiCPlus”. In: Nature communications 9.1, p. 750.

Zhu, Hongyu et al. (2021). “Attention mechanisms in CNN-based single image super-resolution: A brief review and a new perspective”. In: Electronics 10.10, p. 1187.

