## Supplemental Document for "ScHiCAtt: Enhancing Single-Cell Hi-C Resolution Using Attention-Based Models"


**Table 1.** Performance Metrics for ScHiCAtt with Different Loss Function Configurations. The table shows the weights assigned to each loss component and the resulting performance metrics. The optimal configuration is highlighted in bold.

| **Configuration** | $\alpha$ (MSE) | $\beta$ (Perceptual Loss) | $\gamma$ (TV Loss) | $\delta$ (Adversarial Loss) | **PSNR** | **SSIM** | **SNR** |
|---|---|---|---|---|---|---|---|
| 1 | 0.6 | 0.2 | 0.1 | 0.1 | 38.50 | 0.9750 | 5400.25 |
| 2 | 0.5 | 0.4 | 0.05 | 0.05 | 39.10 | 0.9800 | 5450.30 |
| 3 | 0.4 | 0.3 | 0.2 | 0.1 | 37.80 | 0.9700 | 5350.10 |
| 4 | 0.5 | 0.3 | 0.1 | 0.1 | **40.00** | **0.9835** | **5550.75** |
| 5 | 0.6 | 0.1 | 0.2 | 0.1 | 37.20 | 0.9650 | 5300.00 |
| 6 | 0.5 | 0.2 | 0.15 | 0.15 | 39.50 | 0.9810 | 5480.60 |

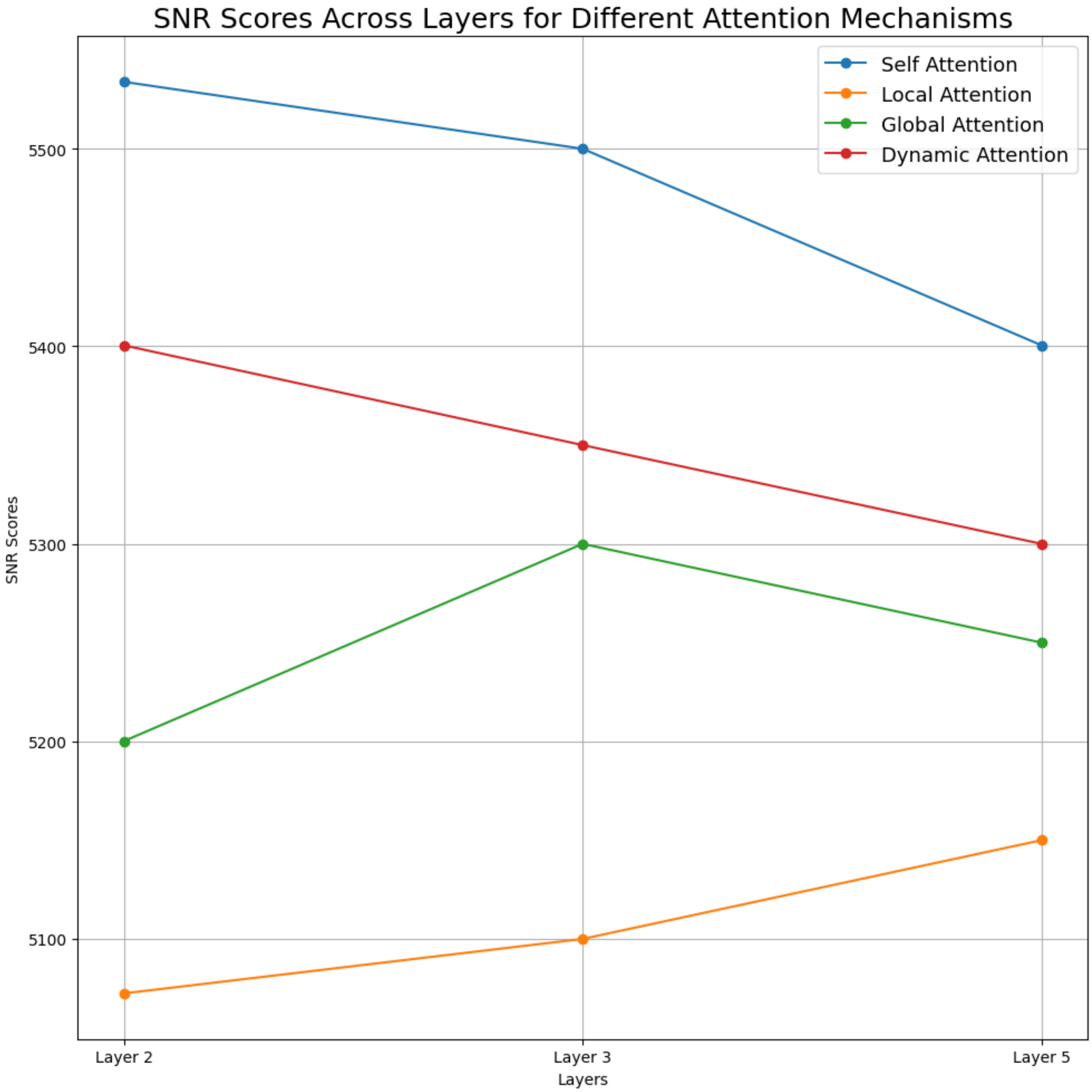


**Fig. 1.** Benchmarking SNR scores across layers for different attention mechanisms on the Human Cell 1 dataset. These are the average scores derived from the analysis of all chromosomes under investigation.

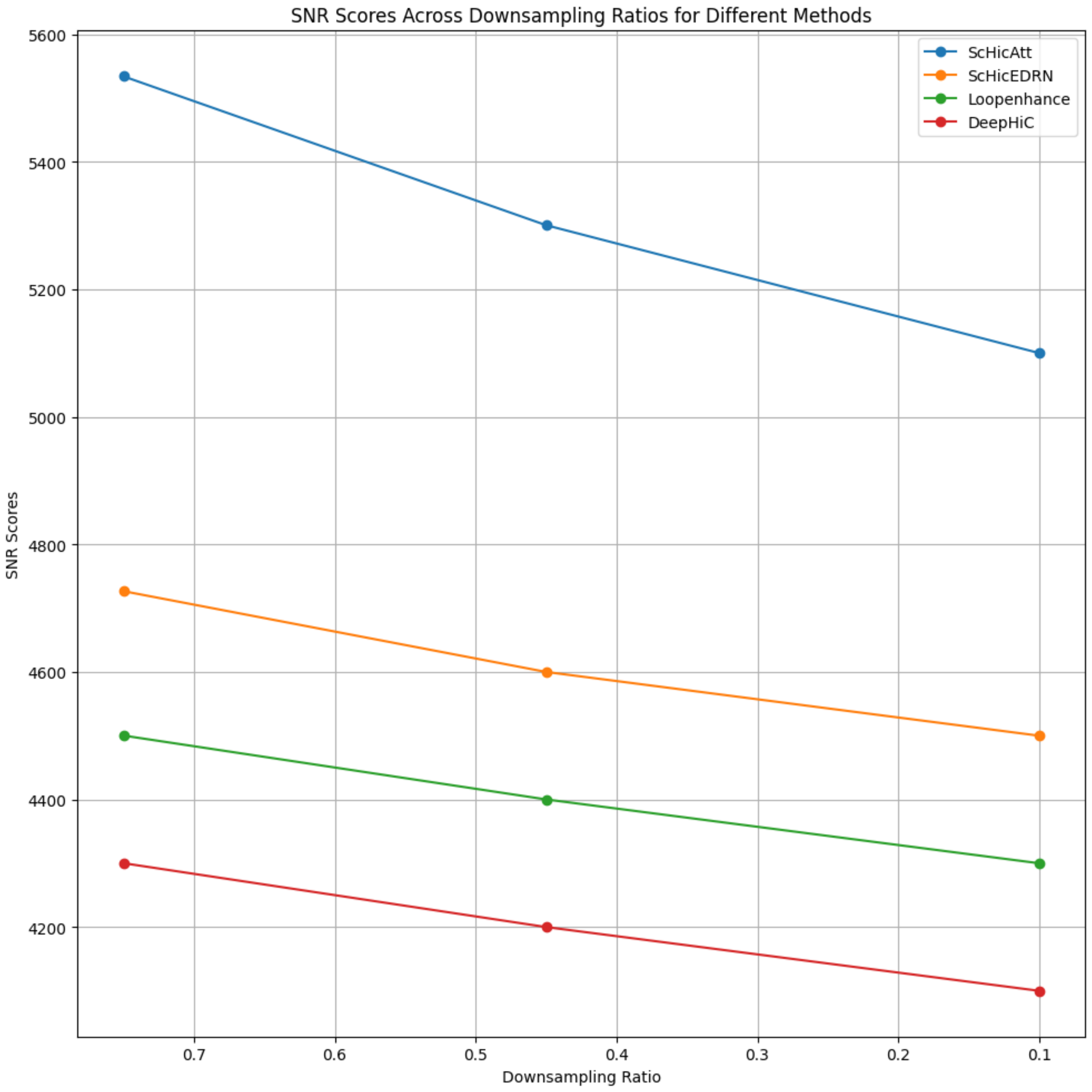


**Fig. 2.** Benchmarking SNR scores of ScHiCAtt and other algorithms across Downsampling Ratio on the Human Cell 1 dataset. These are the average scores derived from the analysis of all chromosomes under investigation.

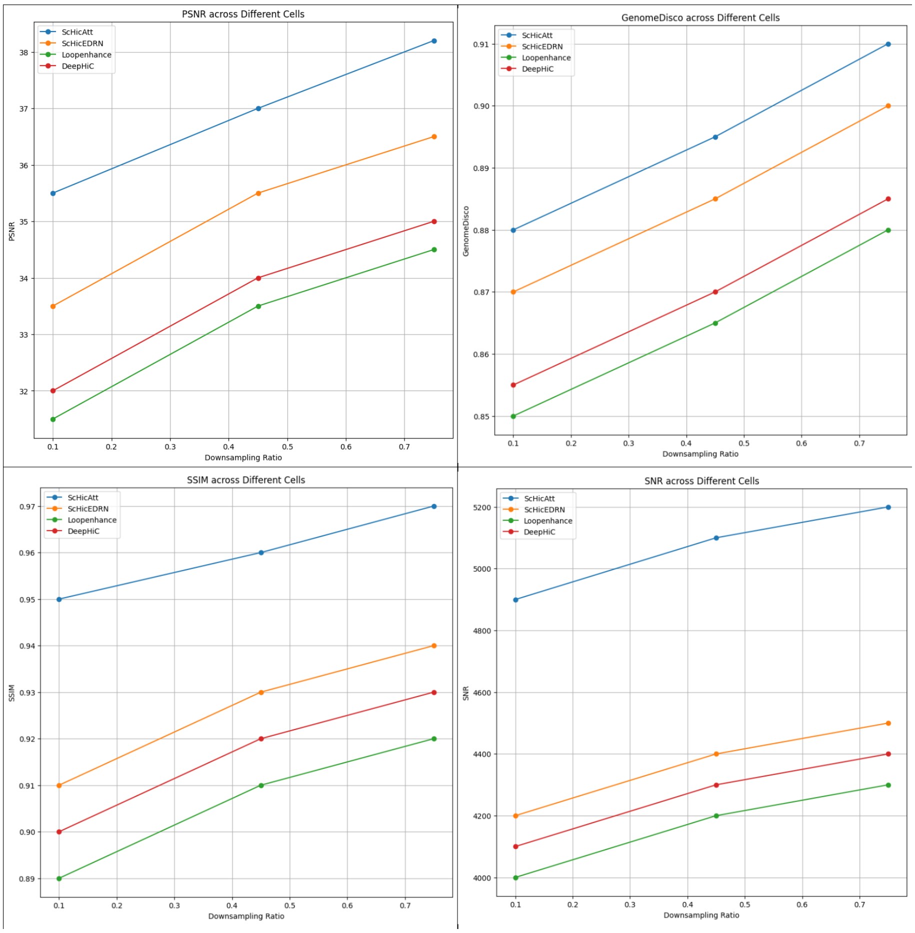


**Fig. 3.** Average performance of different methods across various downsampling ratios when trained on one cell and tested on other cells for chromosomes 2,6,10 and 12 of Human Cell 2. ScHiCAtt consistently outperforms other methods across most metrics.

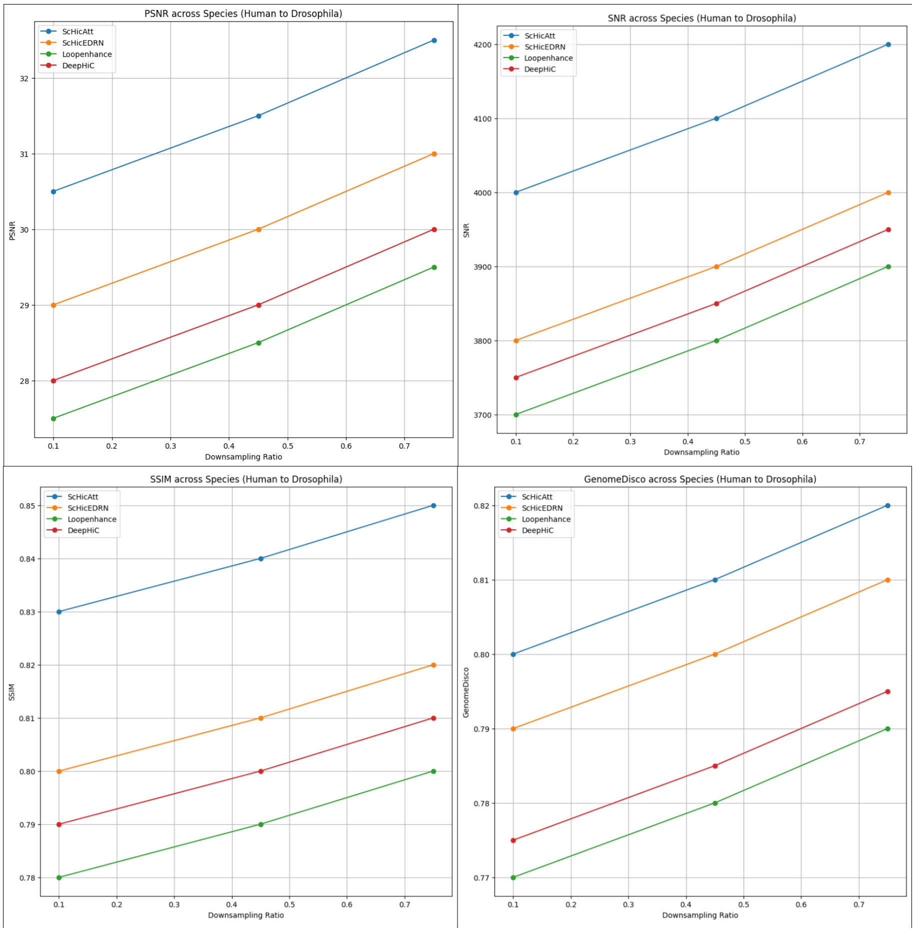


**Fig. 4.** Average performance of different methods across various downsampling ratios when trained on human data and tested on Drosophila data on chromosome X and 2L.ScHiCAtt generally outperforms other methods across most metrics, even in a cross-species scenario.

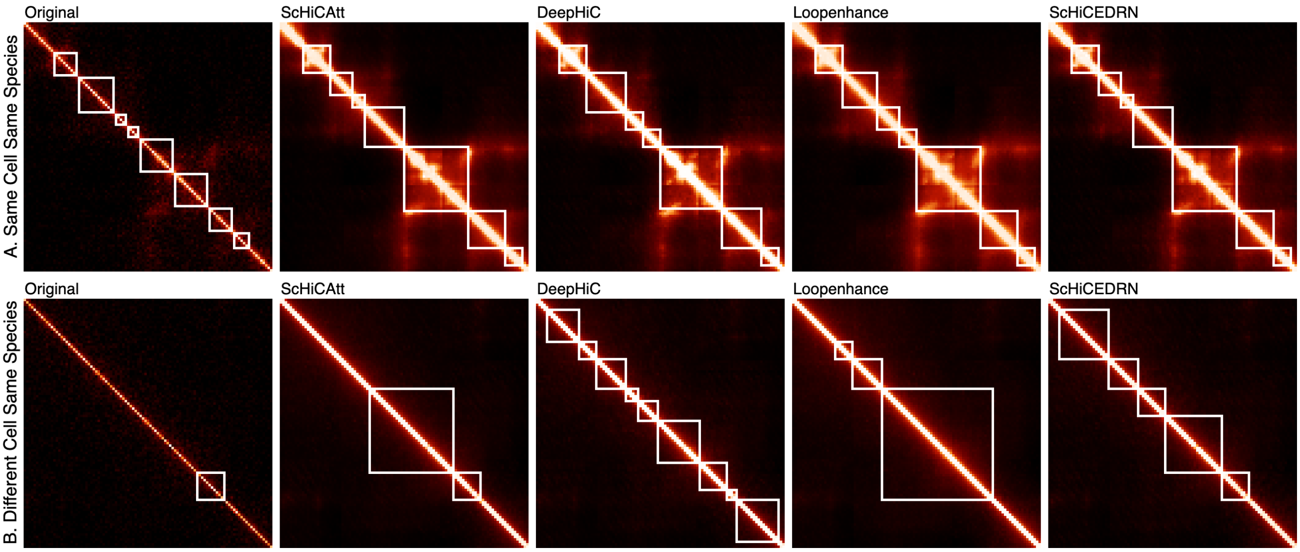


**Fig. 5. Visualization of TAD boundaries.** Visualization of TAD boundaries from 500 to 600 genomic bin regions of A. Same Cell Same Species, B. Different Cell Same Species model generated matrix across original, ScHiCAtt, DeepHiC, Loopenhance, and ScHiCEDRN using Human Cell 2 at $40Kb$ resolution.
